# The level of endothelial glycocalyx maturity modulates interactions with charged nanomaterials

**DOI:** 10.1101/2024.09.10.611831

**Authors:** Claire A. Bridges, Lu Fu, Jonathan Yeow, Xiaojing Huang, Miriam Jackson, Rhiannon P. Kuchel, James D. Sterling, Shenda M. Baker, Megan S. Lord

**Affiliations:** Graduate School of Biomedical Engineering, University of New South Wales, Sydney, NSW 2052, Australia; Molecular Surface Interaction Laboratory, Mark Wainwright Analytical Centre, University of New South Wales, Sydney, NSW 2052, Australia; Electron Microscope Unit, Mark Wainwright Analytical Centre, University of New South Wales, Sydney, NSW 2052, Australia; College of Innovation, Entrepreneurship, and Economic Development, Missouri University of Science and Technology, Rolla, MO 65409, USA; Synedgen Inc, Claremont, CA 91711, USA

**Keywords:** nanoparticle, endothelium, glycocalyx, drug delivery

## Abstract

Nanomaterials have been extensively investigated for their potential in delivering therapeutics to target tissues, but few have advanced to clinical application. The luminal surface of endothelial cells that line blood vessels are covered by a glycocalyx, a complex extracellular matrix rich in anionic glycans. However, the role of this glycocalyx in governing nanomaterial-cell interactions is often overlooked. In this study, we demonstrate that gold nanoparticles functionalized with branched polyethyleneimine (AuNP+) bind to primary human endothelial cells expressing either a developing or mature glycocalyx, with the interaction involving hyaluronan and heparan sulfate. Notably, the mature glycocalyx decreases the toxicity of AuNP+. In contrast, lipoic acid-functionalized gold nanoparticles (AuNP-) bind to endothelial cells with a developing glycocalyx, but not a mature glycocalyx. To further investigate this phenomenon, we studied charged polymers, including poly(arginine) (polyR) and poly(glutamic acid) (polyE). PolyE does not associate with endothelial cells regardless of glycocalyx maturity, but when glycans are enzymatically degraded, it can bind to the cells. Conversely, polyR associates with endothelial cells irrespective of glycocalyx maturity or glycan degradation. These findings highlight the intricate relationship between nanomaterial charge and presentation in interactions with endothelial cells, offering insights for modulating nanomaterial interactions with the blood vessel wall.

## 1. INTRODUCTION

The emergence of engineered nanomaterials has led to considerable anticipation regarding their potential to drive advancements for a wide range of biomedical applications including drug delivery, and diagnostic imaging [1, 2]. Key to the biological performance of engineered nanomaterials is their binding to cell surfaces [3]. Numerous physicochemical properties of engineered nanomaterials, including size, shape, hydrophobicity, and surface charge and functional ligands, have been investigated for their effect on binding to cells and subsequent internalization processes [3, 4]. However, few engineered nanomaterials have successfully progressed from the laboratory to the clinic which is in part due to the disparity in their biological evaluation between *in vitro* and *in vivo* systems [3, 5].

With engineered nanomaterial delivery systems largely intended to be administered via intravenous injection, inhalation, or ingestion this necessitates their interaction with the bloodstream on their path to target cells [6]. Additionally, the endothelium is an important target for therapy being involved in numerous pathophysiologic conditions including atherosclerosis, tumor growth, and restenosis [7]. Thus, the ability of engineered nanomaterials to bind and transit endothelial cells, either by paracellular or transcellular mechanisms, lining the blood vessel wall is a key consideration in determining their biological fate [6]. Underappreciated, yet critical in regulating transport to and across all mammalian cell membranes, is the glycocalyx which extends several hundred nanometres from the cell membrane [8, 9]. The endothelial glycocalyx is crucial for functions of the endothelium [10, 11]. Remarkably, however, the glycocalyx has largely been overlooked in the biological assessment of engineered nanomaterials [9].

With the glycocalyx taking a prominent position at the endothelial cell surface it is the first point of interaction between engineered nanomaterials and cells. Critically, we are yet to fully understand the impact of the glycocalyx on interactions with engineered nanomaterials. Such interactions will likely depend on the physicochemical properties of the engineered nanomaterials and their ability to interact with the glycocalyx, which behaves like an anionic hydrogel [9]. This interaction has so far been shown to be electrostatic in nature like other biological interactions with glycans [12] which is in contrast to the receptor-mediated binding reported at the cell membrane [13].

The present study explores the interaction of gold nanoparticles (AuNP), which are widely studied for their broad biomedical applications [14] when functionalized with either cationic or anionic ligands to examine the role of surface charge in binding to the endothelial glycocalyx and subsequent intracellular uptake. Building on previous research that demonstrated the exclusion of small (10 nm), charged PEG functionalized AuNPs by the endothelial glycocalyx both *in vitro* and *in vivo* [15-17], we focus on the effects of nanoparticle charge. Unlike most studies that assess nanoparticle performance in short-term *in vitro* cultures, we culture primary human endothelial cells over a time course that allows for glycocalyx formation and VE-cadherin junction development, mimicking functional endothelium *in vivo* [9, 18]. We show that 50 nm cationic branched polyethyleneimine (PEI) functionalized AuNPs (AuNP+) associate with endothelial cells regardless of whether the glycocalyx is developing or mature. In contrast, 50 nm lipoic acid functionalized AuNPs (AuNP-) bind to cells with a developing glycocalyx but are excluded by those with a mature glycocalyx. This charge-dependent exclusion is also observed with charged polymers, such as poly(arginine) and poly(glutamic acid), although polymer interactions with the glycocalyx do not depend on its maturity.

## 2. MATERIALS AND METHODS

All reagents were purchased from Sigma-Aldrich unless stated otherwise.

### 2.1. Culture of endothelial cells to express a glycocalyx

Primary human umbilical vein endothelial cells (HUVECs, Lonza) were maintained in endothelial cell growth medium (EGM™-2 BulletKit, Lonza) at 37 °C with 5% CO_2_ and used at passages 2 - 8. The medium was replaced every 2 to 3 days. HUVECs were seeded in tissue culture plates at a density of 1,429 cells/cm^2^ in medium and cultured for up to 7 days prior to use in experiments.

### 2.2. Assays to characterize the endothelial glycocalyx

#### 2.2.1. Transmission electron microscopy (TEM)

Cells were cultured for 7 days as detailed in section 2.1 and harvested using EDTA (50 mM) in PBS, and pelleted. Cells were then fixed for 30 min at 22 °C in Karnovski’s Fixative (paraformaldehyde (2.5% w/v), glutaraldehyde (2% v/v) in Sorensen’s phosphate buffer). Fixed cells were embedded in agar (2% w/v) and then stained using a BiowavePro+ (Pelco) with osmium tetroxide (1% w/v) for 10 min followed by uranyl acetate (2% w/v) for 2 min at 22 °C, under vacuum, followed by dehydration through a graded ethanol series and finally embedded in Pure LR White resin. Samples were polymerized at 60 °C for 24 h. Ultrathin sections were cut via ultramicrotomy and post-stained with uranyl acetate (2% w/v, 20 min) and lead citrate (0.5% w/v, 5 min) prior to washing and imaging with a transmission electron microscope (JEM-1400, JEOL).

#### 2.2.2. Confocal microscopy

Cells were cultured in 18-well μ-slides (ibidi GmbH) and incubated with wheatgerm agglutinin (WGA) conjugated with Alexa Fluor 647 for glycans containing sialic acid (5 μg/mL diluted in medium) for 10 min at 37 °C followed by two washes with PBS and then fixed in paraformaldehyde (4% w/v) in PBS for 15 min at 37 °C in the dark. Alternatively, cells were permeabilized with sucrose (300 mM), NaCl (50 mM), MgCl_2_ (3 mM), HEPES (2 mM), and Triton X-100 (0.5% v/v), pH 7.2 for 5 min at 4 °C. The cells were then washed three times in PBS and blocked with goat serum (1% v/v) in PBST (PBS with Tween-20 (0.05% w/v); blocking solution) at room temperature (RT) for 1 h. Cells were then incubated with anti-heparan sulfate (HS) (2 μg/mL, clone 10E4, Amsbio), anti-syndecan-1 (2 μg/mL, clone DL-101, Santa Cruz Biotechnology), anti-CD44 (2 μg/mL, clone Hermes-1, Developmental Studies Hybridoma Bank), or anti-VE-cadherin (1 μg/mL, Abcam) antibodies or biotinylated hyaluronan (HA) binding protein (5 μg/ml, Sigma-Aldrich) diluted in the blocking solution for 16 h at 4 °C then washed twice with PBST. Wells were then incubated with the appropriate Alexa Fluor 488 conjugated secondary antibodies (goat anti-rat IgG (H+L) (4 μg/mL), goat anti-mouse IgG & lgM (H+L) (4 μg/mL), or goat anti-mouse IgG (H+L) (4 μg/mL) (Thermo Fisher Scientific)) or Alexa Fluor 488 conjugated streptavidin (4 μg/mL) for 2 h at RT and then rinsed twice with PBST. Samples were incubated with Hoechst-33342 (1:1000, Thermo Fisher Scientific) for 30 min at RT followed by two washes with PBST and a final rinse with PBS at 4 °C prior to imaging using a confocal microscope (LSM 880, Zeiss) with a Plan-Apochromat 63.0× 1.4 Oil objective.

### 2.3. Nanoparticle and polymer functionalization

AuNPs (50 nm) functionalized with branched polyethyleneimine or lipoic acid (NanoComposix) were used as supplied. Poly(L-arginine)-azide (polyR-azide) (29 kDa; Alamanda Polymers) and poly(L-glutamic acid)-azide (polyE-azide) (26 kDa; Alamanda Polymers) were labelled with Alexa Fluor 488-dibenzocyclooctyne (Jena Bioscience) and purified by extensive dialysis with a Slide-A-Lyzer dialysis casette (Thermo Fisher Scientific) against deionized water as the running buffer. ATR-FTIR (Spotlight 400, Perkin Elmer) was used to measure changes in the surface chemical structure of the polymers following conjugation.

### 2.4. Assays to study nanoparticle or polymer interactions with cells

#### 2.4.1. Cytocompatibility

Cells were cultured as detailed in section 2.1 in 96-well tissue culture plates for 1, 3 or 7 days prior to the addition of AuNP (0-200 μg/mL), poly E (0-24 μg/mL) or polyR (0-24 μg/mL) to the medium for 2 h, or medium alone (positive control). The CyQuant assay was performed as per the manufacturer’s instructions. Selected cells were treated with heparinase III (20 mU/mL; Ibex Pharmaceuticals), hyaluronidase from bovine testes (20 U/mL) and neuraminidase from *Clostridium perfringens* (*C. welchii*) (60 mU/mL) prior to AuNP or polymer exposure.

#### 2.4.2. Microscopy

Cells were cultured as detailed in section 2.1 in 18-well μ-slides (ibidi GmbH) for either 1, 3 or 7 days prior to incubation with AuNPs (3 μg/mL), polyE (0-24 μg/mL) or polyR (3 μg/mL) for 2 h. The cells were washed twice with PBS before being incubated with antibodies or probes for glycocalyx components as detailed in section 2.2.2 or CellMask™ Orange plasma membrane stain (1:1000; Thermo Fisher Scientific) diluted in culture medium for 10 min at 37 °C. Samples were then washed twice with PBS and then fixed in paraformaldehyde (4% w/v) in PBS for 15 min at 37 °C in the dark. Samples were incubated with Hoechst-33342 (1:1000 in blocking solution, Thermo Fisher Scientific) for 30 min at RT followed by two washes with PBST and a final rinse with PBS at 4 °C prior to imaging using a confocal microscope (LSM 880, Zeiss) with a Plan-Apochromat 63.0× 1.4 Oil objective. AuNP were visualized utilizing the multi-photon imaging capabilities of the confocal microscope using a Tunable InSight DeepSee laser (800 nm, MKS/Spectra-Physics) with BiG non-descanned GaAsP detectors (Zeiss). Co-localization analysis was performed using the Manders’ colocalization coefficient (MCC) analysis in Fiji (ImageJ) with the “Colocalisation” plugin from the Genome Damage and Stability Centre (GDSC) at the University of Sussex. Volumetric analysis of AuNP, polyE, and polyR was performed with Imaris 9.9.1 imaging software (Oxford Instruments).

#### 2.4.3. Flow cytometry

Cells were cultured as detailed in section 2.1 in 12-well tissue culture plates for 1, 3 or 7 days and then incubated with AuNPs (3 μg/mL), polyE (3 μg/mL) or polyR (3 μg/mL) for 2 h at 37 °C. Selected cells were treated with heparinase III, hyaluronidase, and neuraminidase as detailed in section 2.4.1 prior to polymer exposure. The cells were then washed twice with PBS before being incubated with Zombie NIR Fixable Viability dye (1:1000, BioLegend) for 15 min at RT, protected from light. Cells were then washed three times with flow buffer before being fixed in paraformaldehyde (4% w/v) in PBS for 15 min at 37 °C in the dark. Cells were then washed three times and resuspended in flow buffer (150 μL). Samples were stored at 4 °C, protected from light, until analysis in a flow cytometer (FACSymphony™ A3, BD) for fluorescence intensity, forward scatter (FSC) and side scatter (SSC) for a minimum of 10^4^ events after gating for live, single cells. Data were analyzed with the FlowJo_V10 software (BD).

### 2.5. Assay to study AuNP interactions with isolated glycans by quartz crystal microbalance with dissipation (QCM-D) monitoring

A continuous flow of 50 μL/min and temperature of 37 °C were applied throughout the experiments using a QCM-D (Analyzer, Q-Sense). After a baseline was established with deionized water (filtered and degassed), the sensor surfaces were coated with either heparin thiol (50 μg/mL; 27 kDa; Creative PEGWorks) or hyaluronan thiol (50 μg/mL; 50 kDa; Creative PEGWorks) for 10 min, washed with deionized water, blocked with thiolated PEG (50 μg/mL; 500 Da; Creative PEGWorks), washed with deionized water, and then exposed to AuNP+ or AuNP-(50 μg/mL) followed by a deionized water rinse. Binding at each step was monitored by changes in frequency (Δ*f*) and dissipation (ΔD) at the fundamental frequency as well as the 3^rd^ –11^th^ overtones.

### 2.6. Statistical analyses

Statistically significant differences were determined by one-way analysis of variance (ANOVA) and the Tukey post-test after assessment of data normality using Prism version 10 (GraphPad). Data are expressed as mean ± standard deviation (SD) unless stated otherwise. Statistical significance was accepted at *p*≤0.05 and indicated in the figures as \**p*≤0.05, \*\**p*≤0.01, \*\*\**p*≤0.001, and \*\*\*\**p*≤0.0001.

## 3. RESULTS

### 3.1 Cationic nanoparticles associate with endothelial cells to a greater extent than anionic nanoparticles

We first explored the interaction of charged nanoparticles with endothelial cells expressing a glycocalyx. We examined AuNP functionalized with either branched polyethyleneimine (AuNP+) or lipoic acid (AuNP-), representing cationic and anionic nanoparticles, respectively (**Figure 1A**). Endothelial cells were cultured for 7 days to develop a mature glycocalyx, which is estimated to extend 350 nm from the cell membrane, based on TEM imaging (**Figure 1B**). To assess AuNP interactions with these cells, a cytotoxicity assay was conducted to determine the appropriate dose range for further analysis. AuNP-are cytocompatible up to 100 μg/mL (**Figure 1C**), while AuNP+ exhibit cytotoxicity at doses above 25 μg/mL (**Figure 1D**). Quantitative analysis of confocal microscopy images reveals that AuNP+ associate with endothelial cells 7 times more (*p*<0.0001) than AuNP-after 2 h of exposure (**Figure 1E-F**). Furthermore, AuNP-show no significant association with endothelial cells compared to the control group without exposure to AuNPs (*p*>0.05) (**Figure 1E-F**), indicating negligible interaction between AuNP- and cells cultured for 7 days to express a mature glycocalyx.

**Figure 1.**
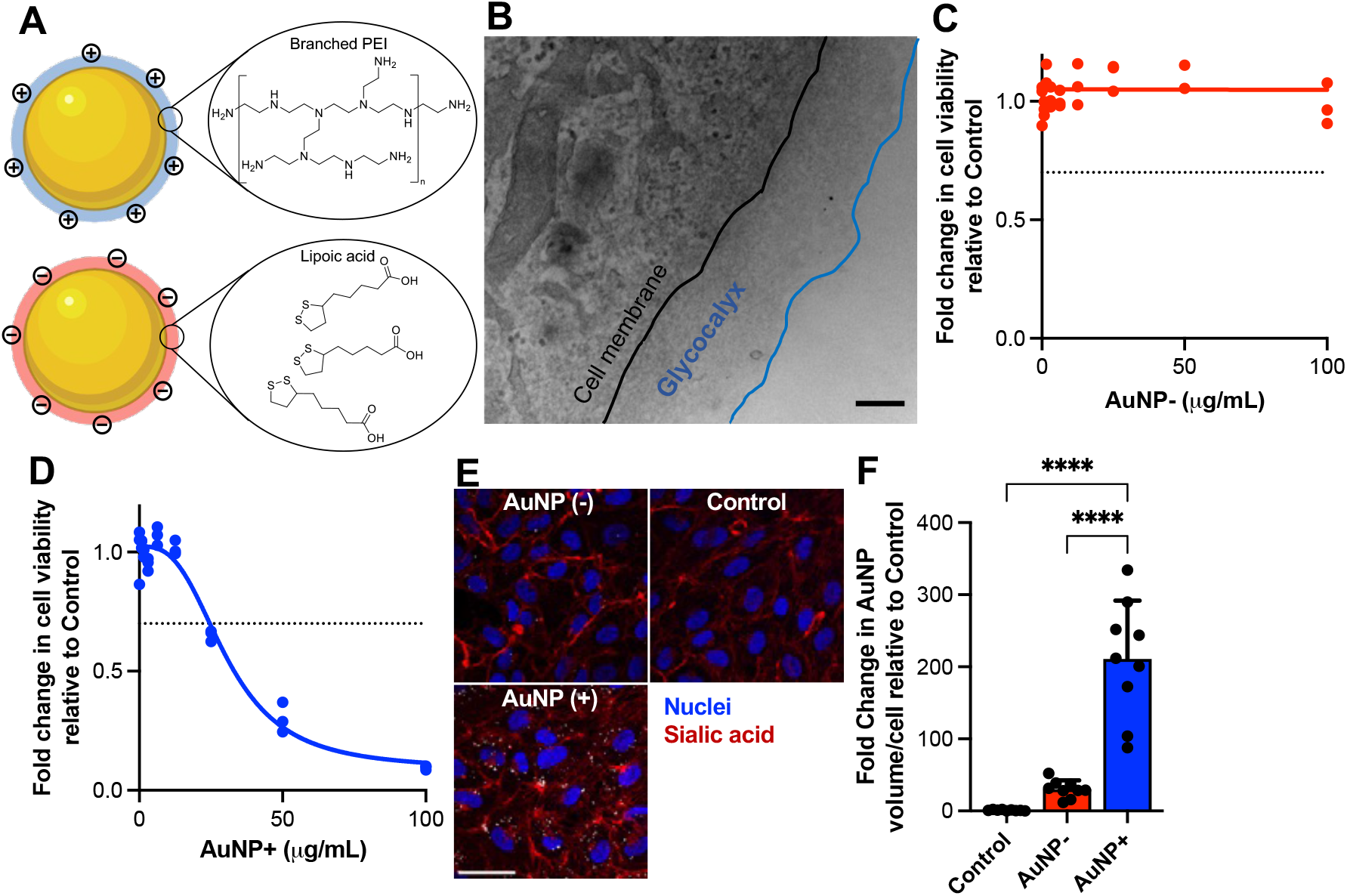
Cationic gold nanoparticles (AuNP+) associate with endothelial cells expressing a glycocalyx. **(A)** Schematic of the structure of AuNPs functionalized with branched polyethyleneimine (AuNP+) or lipoic acid (AuNP-). **(B)** Representative TEM image of endothelial cells following 7 days in culture showing the location of the glycocalyx which extends approximately 350 nm from the cell membrane. Scale bar is 100 nm. Cytotoxicity of **(C)** AuNP- and **(D)** AuNP+ to endothelial cells cultured for 7 days prior to exposure to AuNPs for 2 h over the dose range of 0-100 μg/mL determined by the CyQuant assay. Data are mean ± SD (n=3). **(E)** Representative confocal microscopy images show the association of either AuNP+ or AuNP-(white; 3 μg/mL) with endothelial cells. Cells were counterstained for the cell surface glycans (WGA; red) and nuclei (Hoechst-33342). Scale bar is 40 μm. **(F)** Volumetric measure of AuNP+ or AuNP-signal (white pixels) per cell as a measure of cell association. Data are mean ± SD (n=9). \*\*\*\**p*≤0.0001.

Given the association of AuNP+ with endothelial cells, it was of interest to examine whether AuNPs bind to components of the glycocalyx. The binding of AuNP with the major glycocalyx glycans, hyaluronan (HA) and heparin as a model for heparan sulfate (HS), was analyzed using the QCM-D. For these experiments, gold QCM-D sensors were first coated with either heparin thiol or HA thiol, then the remaining gold sensor surface was blocked with PEG thiol, and lastly exposed to AuNPs (**Figure 2A**). Binding of each of these molecules/AuNPs to the sensor surface is measured by a change in frequency (Δ*f*), which is related to the amount of bound mass, and a change in dissipation (ΔD), which is related to the viscoelasticity of the immobilized molecules (**Figure 2A**). Comparison of the binding of AuNP to heparin thiol was analyzed by the Δ*f* for AuNP binding normalized to the Δ*f* for heparin thiol binding (**Figure 2B**). This analysis indicates that AuNP+ bind to heparin while AuNP-do not (*p*<0.001) (**Figure 2B**). Similarly, AuNP+ bind to HA while AuNP-do not (*p*<0.01) (**Figure 2C**). Comparison of the AuNP+ adsorption behavior to heparin and HA was performed by plotting ΔD versus Δ*f* (D*f* plots; **Figure 2D-E**). These plots reveal that AuNP+ binding to heparin leads to decreases in both Δ*f* and ΔD, indicating that the binding results in a stiffer layer on the sensor surface, consistent with displacement of the hydration shell of heparin due to AuNP+ binding through a condensation mechanism. In contrast, AuNP+ binding to HA results in an initial phase where there are decreases in Δ*f* and increases in ΔD suggesting binding without condensation followed by a second phase where there are decreases in both Δ*f* and ΔD suggesting binding with condensation. Together, these data indicate that AuNP+ bind to major glycocalyx glycans, HA and HS, while AuNP-do not. Furthermore, there are differences in the mechanisms of AuNP+ binding to HA and HS.

**Figure 2.**
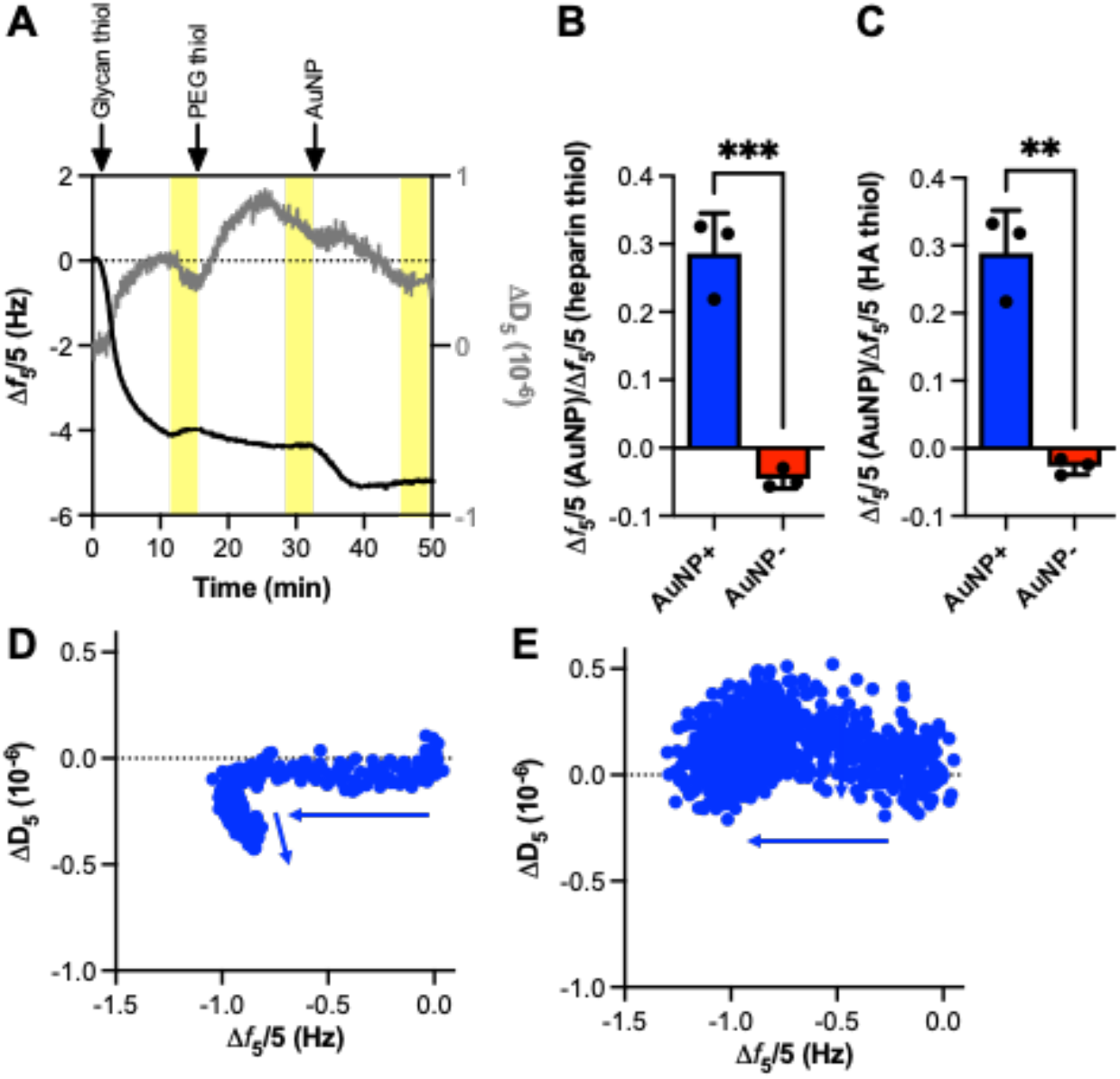
Cationic gold nanoparticles (AuNP+) bind to immobilized heparin and HA. **(A)** Representative QCM-D experiment of AuNP+ binding to immobilized heparin thiol with the sensor surface blocked with PEG thiol (500 Da) prior to addition of AuNP using deionized water as the running buffer displayed as changes in frequency (Δ*f*) and dissipation (ΔD) versus time for the 5^th^ overtone. The regions of the graph highlighted in yellow indicate the addition of running buffer and the arrows indicate the addition of (1) heparin thiol (50 μg/mL), (2) PEG thiol (50 μg/mL), and (3) AuNP (50 μg/mL). **(B)** Relative binding of AuNP to immobilized heparin thiol presented as Δ*f* for AuNP binding divided by Δ*f* for heparin thiol (5^th^ overtone) (n=3). **(C)** Relative binding of AuNP to immobilized HA thiol presented as Δ*f* for AuNP binding divided by Δ*f* for hyaluronan thiol (5^th^ overtone) (n=3). (Representative ΔD versus Δ*f* plot (D*f* plot) for AuNP+ binding to (**D)** heparin thiol, and **(E)** HA thiol. The arrow indicates the time course of the data points. \*\**p*≤0.01 and \*\*\**p*≤0.001.

Having established that AuNP+ bind to HA and HS in an *in vitro* system using isolated molecules, we next determined whether AuNP+ bind to these glycans and other glycocalyx components on the surface of endothelial cells. Confocal microscopy imaging indicates that AuNP+ associate with the plasma membrane (**Figure 3A**) as well as glycocalyx components including syndecan-1, a transmembrane protein decorated with HS, and CD44, a transmembrane protein and principal HA receptor (**Figure 3B**). Additionally, AuNP+ colocalize with multiple glycans in the glycocalyx including, HS and HA (**Figure 3C**). Manders’ colocalization coefficient analysis also indicates that the AuNP+ do not bind to all available glycocalyx or plasma membrane epitopes (**Figure 3D**). Together, these data show that AuNP+ bind to multiple glycocalyx components enabling association with endothelial cells.

**Figure 3.**
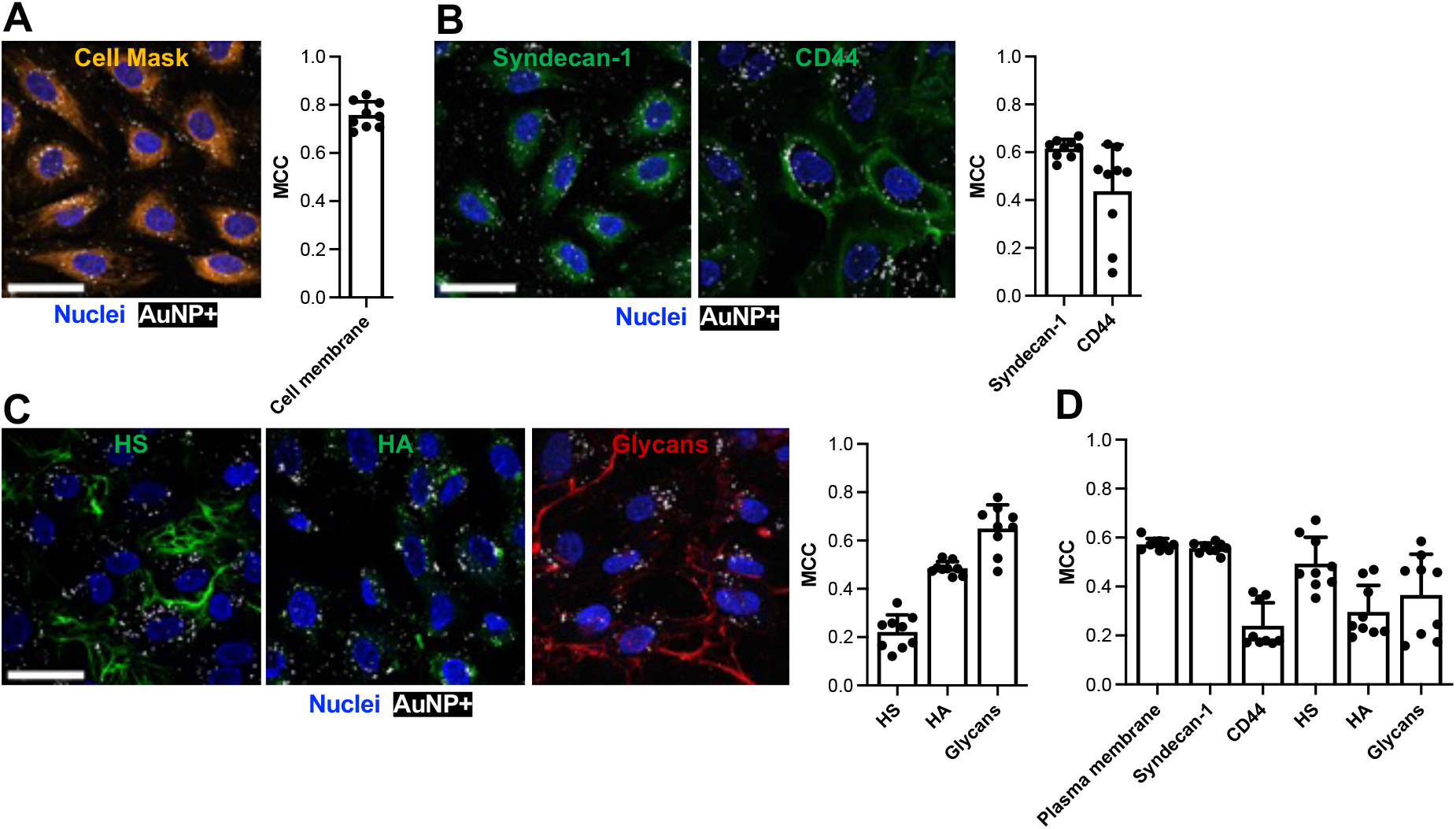
AuNP+ colocalizes with multiple glycocalyx components at the endothelial cell surface. **(A)** Representative confocal microscope image of endothelial cells exposed to AuNP+ (3 μg/mL; white) for 2 h and counterstained for the **(A)** plasma membrane (red; CellMask™), **(B)** syndecan-1 (green; antibody clone DL-101), and CD44 (green; antibody clone Hermes-1) and **(C)** HS (green; antibody clone 10E4), HA (green; HA binding protein) and glycans (WGA; red) and nuclei (blue; Hoechst-33342). Scale bar is 40 μm. Manders’ colocalization coefficient (MCC) analysis for the extent of AuNP+ (white pixels) colocalized with **(A)** cell membrane, **(B)** glycocalyx proteins or **(C)** glycans. Data are mean ± SD (n=9).

Having established that AuNP+ bind to the glycocalyx expressed by endothelial cells after 7 days in culture, a protocol we previously established as optimal for both robust glycocalyx expression and VE-cadherin expression representing functional endothelium [18], we next sought to understand the development of the glycocalyx over this period. The expression of glycocalyx components, including CD44, syndecan-1, HS, HA and glycans significantly increased with increasing culture time (**Figure 4A-B**). Additionally, using a higher cell density than we previously reported [18], the cells expressed VE-cadherin containing cell-cell junctions throughout the measurement period (**Figure 4A**). Collectively, these data indicate that cells require time in culture to establish a mature glycocalyx, irrespective of the expression of VE-cadherin containing cell-cell junctions (**Figure 4C**).

**Figure 4.**
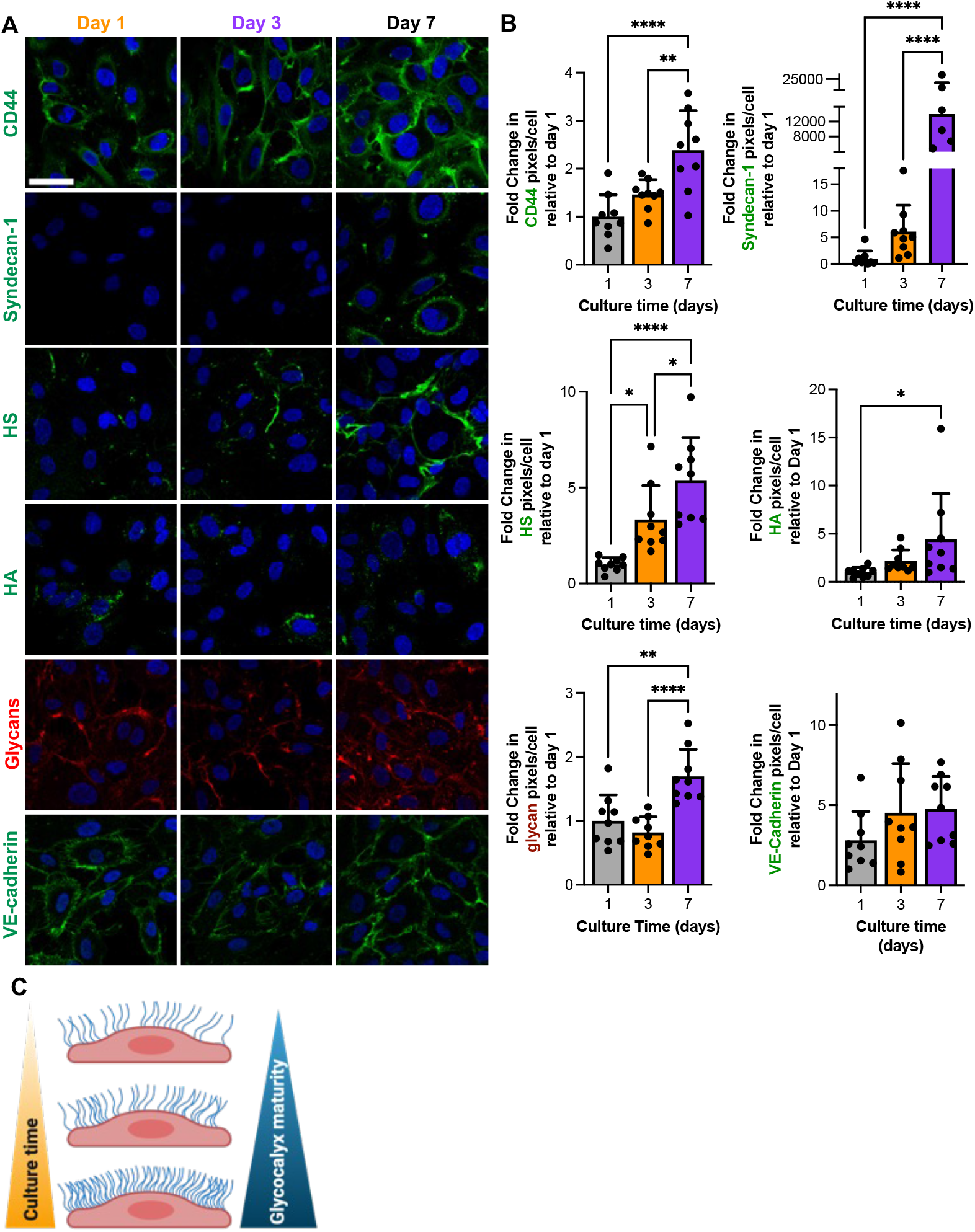
The endothelial glycocalyx matures with increasing time in culture. **(A)** Representative confocal microscopy images show expression of syndecan-1, CD44, HS, HA (green), glycans (red; WGA) and VE-cadherin (green) by endothelial cells after 1, 3 or 7 days of culture. Cells were counterstained for nuclei (blue; Hoechst-33342). Scale bar is 40 µm. **(B)** Quantification of the fold change in glycocalyx component pixels/cell relative to day 1. Data are mean ± SD (n=6-9). \**p*≤0.05, \*\**p*≤0.01 and \*\*\*\**p*≤0.0001. **(C)** Schematic of the maturation of the endothelial glycocalyx with increasing time in culture.

Given that the glycocalyx develops over time in culture, we aimed to investigate how its maturity influences interactions with AuNP. Our results show that AuNP-are cytocompatible across the tested dose range up to 100 μg/mL, regardless of the glycocalyx maturity (**Figure 5A**). In contrast, AuNP+ exhibit cytotoxicity to endothelial cells cultured for 1 or 3 days at doses exceeding 3 μg/mL, which is an 8-fold lower threshold than in cells cultured for 7 days, where a mature glycocalyx is present (**Figure 5B**). These findings suggest that the maturity of the glycocalyx plays a crucial role in modulating the toxicity of AuNP+.

**Figure 5.**
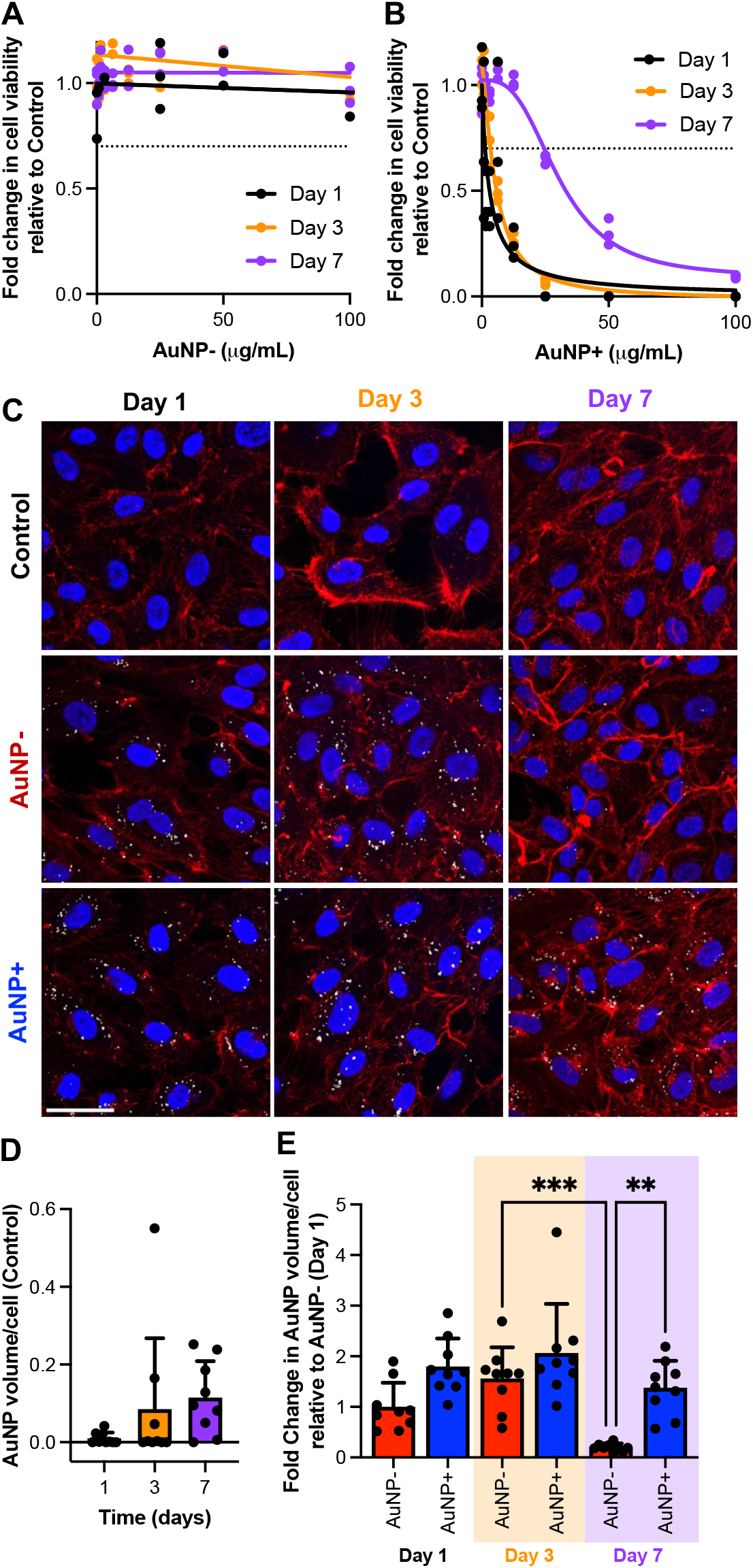
Glycocalyx maturity alters AuNP interactions with endothelial cells. Cytotoxicity of **(A)** AuNP- and **(B)** AuNP+ to endothelial cells cultured for 1, 3 or 7 days prior to exposure to AuNP for 2 h over the dose range of 0-100 μg/mL determined by the CyQuant assay. Data are mean ± SD (n=3). **(C)** Representative confocal microscopy images show association of AuNP+ and AuNP-(3 μg/mL) compared to cells not exposed to AuNP (control) with endothelial cells cultured for 1, 3 or 7 days prior to exposure to AuNPs for 2 h. Cells were counterstained for the cell surface glycans in the glycocalyx (WGA; red) and nuclei (blue; Hoechst-33342). Scale bar is 40 µm. **(D)** Quantification of the volumetric white channel pixel count (background measurement in the Control condition). Data are mean ± SD (n=9). No significant differences measured between conditions. **(E)** Quantification of AuNP association with endothelial cells presented as fold change relative to AuNP-at day 1. Data are mean ± SD (n=9). * *p*≤0.05, ** *p*≤0.01, and *** *p*≤0.001.

Confocal microscopy analysis of the interaction between AuNPs and endothelial cells shows that AuNP-associate similarly with cells cultured for 1 or 3 days, with no significant difference in the extent of association (*p*>0.05) (**Figure 5C-E)**. Although there is no notable difference in the association of AuNP-with cells cultured for 1 versus 7 days, the association is 7.9-fold lower (*p*=0.0002) in cells cultured for 7 versus 3 days (**Figure 5C-E**). In contrast, the association of AuNP+ with endothelial cells remains unaffected by the culture duration (*p*>0.05). Additionally, there is no difference in the association levels of either AuNP-or AuNP+ with cells after 1 or 3 days of culture. However, after 7 days, AuNP+ association is 7-fold higher than that of AuNP-(*p*=0.0014) (**Figure 5C-E**). These results suggest that glycocalyx maturity plays a key role in modulating the interaction of charged nanoparticles with endothelial cells.

With the challenges of fluorescently labelling AuNPs, including strong absorption self-quenching [19], we employed two-photon photoluminescence microscopy for confocal imaging of the nanoparticles. We attempted quantification of AuNP association with cells using flow cytometry, however, at doses below the cytotoxicity threshold, no detectable change in side scatter, a parameter commonly used to quantify metal oxide nanoparticle association with cells [20], was observed (data not shown).

After encountering challenges of quantifying AuNP association with cells using flow cytometry, we redirected our focus to charged polymers, polyE and polyR, which can be more easily labeled for analysis (**Figure 6A**). Similar to anionic AuNPs, polyE is cytocompatible at concentrations analyzed up to 100 μg/mL, regardless of the glycocalyx maturity (**Figure 6B**). In contrast, polyR shows glycocalyx maturity-dependent levels of cytotoxicity levels (**Figure 6C**). Irrespective of endothelial culture time, very little polyE associates with endothelial cells compared to polyR (*p*≤0.001) (**Figure 6D-E**). In contrast, the association of polyR with endothelial cells increases with glycocalyx maturity, with a 1.8-fold higher amount of polyR associating with endothelial cells after 7 days in culture compared to either 1 or 3 days in culture (*p*≤0.05) (**Figure 6D-E**). These results suggest that glycocalyx maturity plays a key role in modulating cationic polymer interactions with endothelial cells.

**Figure 6.**
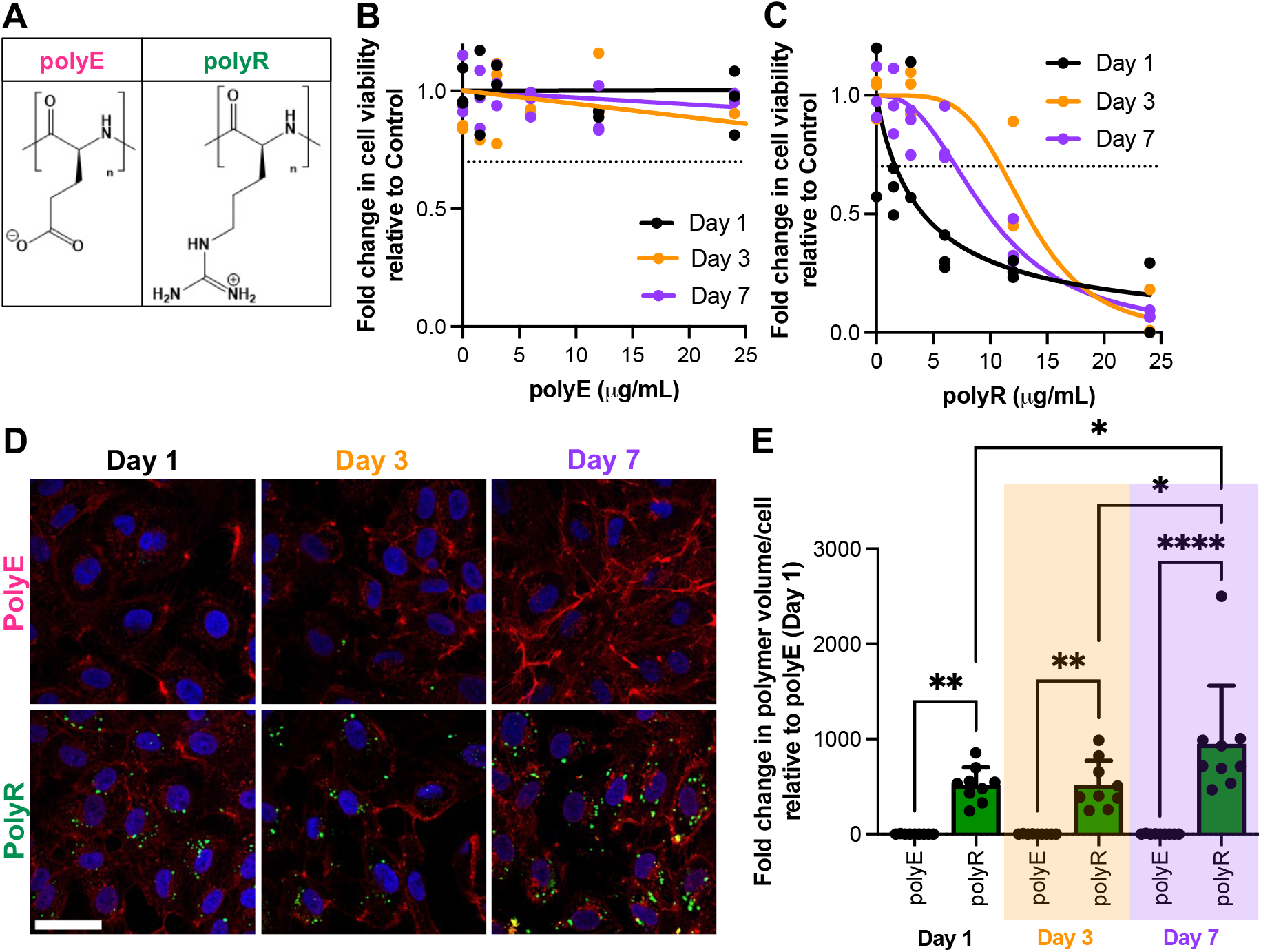
Glycocalyx maturity alters polymer interactions with endothelial cells. **(A)** Schematic of the structure of polyE and polyR. Cytotoxicity of **(B)** polyE and **(C)** polyR to endothelial cells cultured for 1, 3 or 7 days prior to exposure to polymers for 2 h over the dose range of 0-24 μg/mL determined by the CyQuant assay. Data are mean ± SD (n=3). **(D)** Representative confocal microscopy images show association of polymers (3 μg/mL) compared to cells not exposed to polymer (Control) with endothelial cells cultured for 1, 3 or 7 days prior to exposure to polymers for 2 h. Cells were counterstained for the cell surface glycans in the glycocalyx (WGA; red) and nuclei (blue; Hoechst-33342). Scale bar is 40 µm. **(E)** Quantification of polymer association with endothelial cells presented as fold change relative to polyE exposed to endothelial cells after 1 day in culture. Data are mean ± SD (n=9). * *p*≤0.05, ** *p*≤0.01, and *** *p*≤0.001.

To further quantify polyE and polyR interactions with endothelial cells, flow cytometry was employed. Like the confocal microscopy analysis, a negligible level of polyE associates with endothelial cells irrespective of glycocalyx maturity (**Figure 7A**). In contrast, polyR associates with almost all endothelial cells as shown by the shift to increased fluorescence intensity upon exposure to cells after different culture periods (**Figure 7B-C**). In contrast to microscopy, there is no change in the level of association of polyR to endothelial cells with glycocalyx maturity (**Figure 7C**).

**Figure 7.**
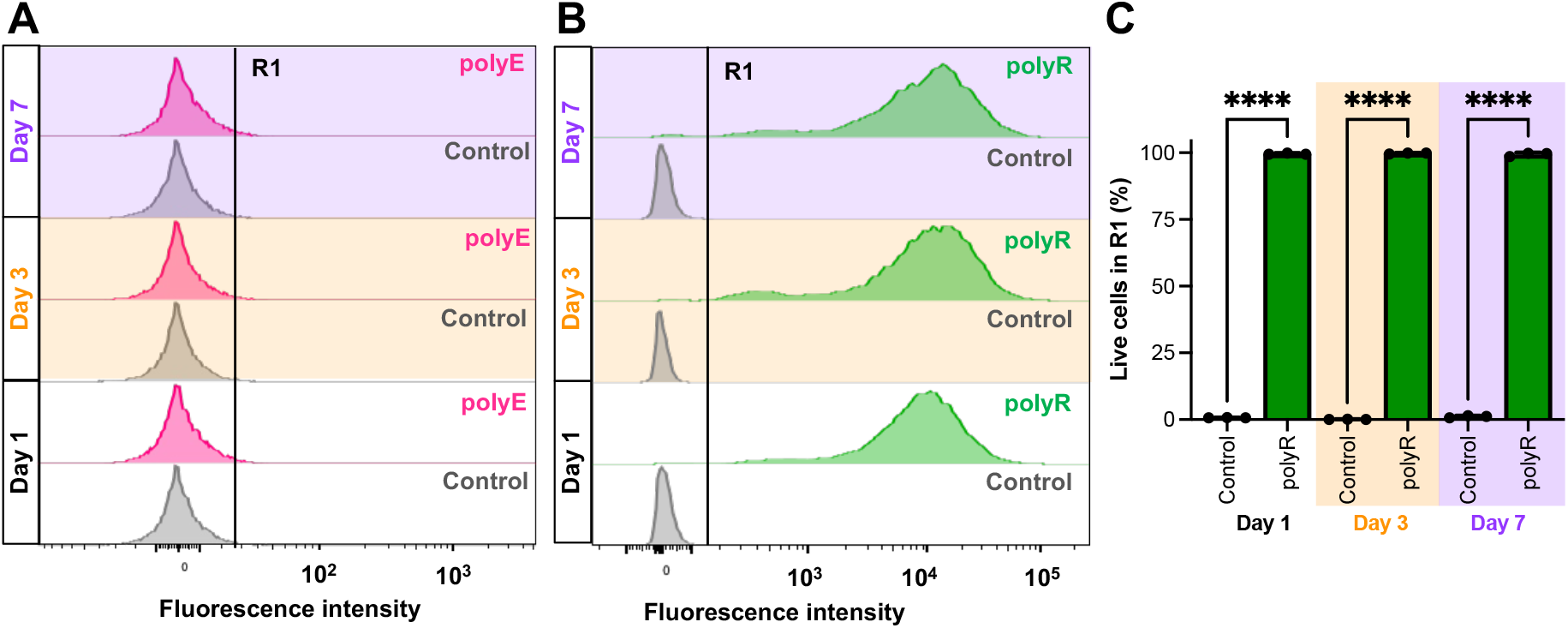
PolyR associates with endothelial cells irrespective of glycocalyx maturity. **(A)** Representative flow cytometry fluorescence intensity profiles of the association of polyE (3 μg/mL) with endothelial cells cultured for 1, 3 or 7 days prior to exposure to polyE for 2 h over compared to cells without exposure to polyE (Control). R1 denotes the region where fluorescence intensity is higher than that of the Control. **(B)** Representative flow cytometry fluorescence intensity profiles of the association of polyR (3 μg/mL) with endothelial cells cultured for 1, 3 or 7 days prior to exposure to polyR for 2 h compared to cells without exposure to polyR (Control). R1 denotes the region where fluorescence intensity is higher than that of the Control. **(C)** Proportion of cells in R1 (defined in panel B). Data are mean ± SD (n=3). **** *p*≤0.0001.

Lastly, to investigate the role of the glycocalyx in modulating the interactions of charged polymers, key glycans were enzymatically degraded using a combination of neuraminidase, which cleaves terminal sialic acid, hyaluronidase, which depolymerizes HA, and heparinase III, which depolymerizes HS, resulting in a degraded glycocalyx (**Figure 8A**). This treatment resembles endothelial glycocalyx degradation during vascular inflammation, such as observed in sepsis and atherosclerosis, where endothelial cell activation facilitates the expression of these glycan degrading enzymes [8, 21]. The cytotoxicity profile of polyE and polyR toward endothelial cells remained consistent, regardless of whether the cells had a mature or degraded glycocalyx (**Figure 8B**). However, the degraded glycocalyx facilitated a 9.4-fold increase in polyE association compared to the mature glycocalyx (*p*<0.0001) (**Figure 8C**). In contrast, polyR association levels were unaffected by the state of the glycocalyx, whether it was mature or degraded (**Figure 8D**). These data confirm the key role of the glycocalyx in modulating biomaterial interactions at the cell surface and the interplay between polymer charge and cell association.

**Figure 8.**
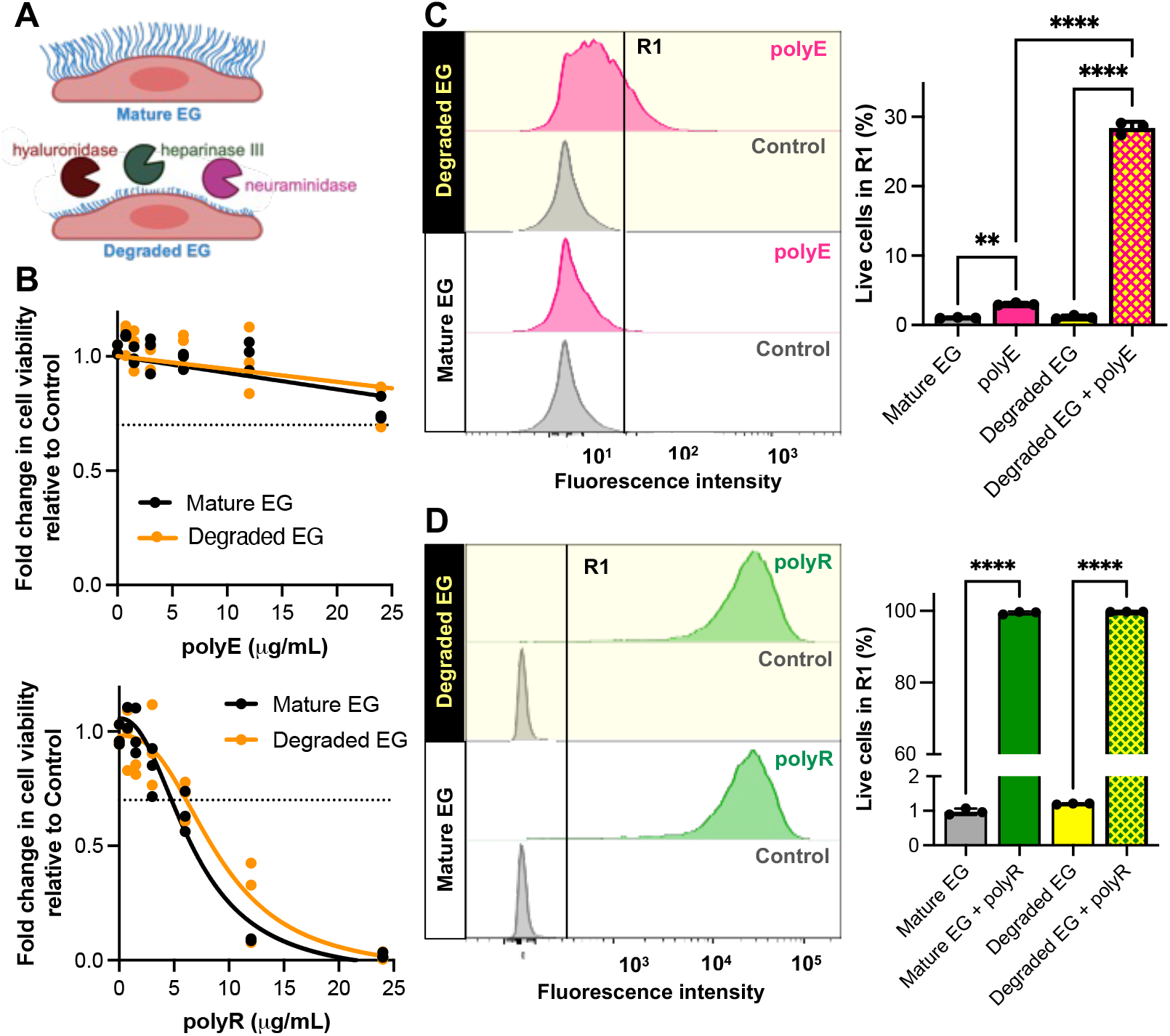
Glycocalyx degradation alters polymer interactions with endothelial cells. **(A)** Schematic of the conditions which were used to measure polymer association with endothelial cells including cells cultured for 7 days to express a mature glycocalyx (Mature EG) and cells cultured for 7 days and then treated with glycan degrading enzymes, hyaluronidase, heparinase III and neuraminidase to produce a degraded glycocalyx (Degraded EG). **(B)** Cytotoxicity of polymers to endothelial cells expressing either a mature or degraded glycocalyx and then exposed to either polymers for 2 h over the dose range of 0-24 μg/mL determined by the CyQuant assay. Data are mean ± SD (n=3). **(C)** Representative flow cytometry fluorescence intensity profiles of the association of polyE (3 μg/mL) with endothelial cells expressing either a mature or degraded glycocalyx and then exposed to polyE for 2 h compared to cells without exposure to polyE (Control). R1 denotes the region where fluorescence intensity is higher than that of the Control. Quantitation of the proportion of cells in R1 (defined in panel C). Data are mean ± SD (n=3). **(D)** Representative flow cytometry fluorescence intensity profiles of the association of polyR (3 μg/mL) with endothelial cells expressing either a mature or degraded glycocalyx and then exposed to polyR for 2 h compared to cells without exposure to polyR (Control). R1 denotes the region where fluorescence intensity is higher than that of the control. Quantitation of the proportion of cells in R1 (defined in panel D). Data are mean ± SD (n=3). ** *p*≤0.01 and **** *p*≤0.0001.

## 4. DISCUSSION

The glycocalyx plays a crucial role in modulating the internalization of molecules in physiological processes, and is increasingly recognized for its influence on the interaction between nanomaterials and cell surfaces [9]. Previous studies have shown that cationic nanomaterials, whether molecular polymers or nanoparticles, can penetrate the glycocalyx and undergo internalization, whereas anionic nanomaterials are generally excluded [18, 22, 23]. In this study, we extend these findings by directly investigating how AuNPs interact with glycans, specifically HA and HS, which are abundant in the endothelial glycocalyx. Our results show that cationic AuNPs (AuNP+) bind to these glycans, while anionic AuNPs (AuNP-) do not. Additionally, we identify distinct mechanisms of AuNP+ binding to HA and HS, indicating varying affinities for different glycans. This sheds light on a possible mechanism by which cationic nanomaterials can bind to and traverse the glycocalyx and be internalized. These findings align with previous research showing that cationic polymers bind to HS [18], and genome-wide genetic screens implicating HS in nanoparticle internalization [24].

Building on our *in vitro* analyses with isolated glycans and nanoparticles, we demonstrate that AuNP+ associate with primary human endothelial cells expressing a mature glycocalyx, while AuNP-do not. This observation aligns with previous *in vitro* and *in vivo* studies, which have shown that various cationic nanoparticles and polymers, such as amine-galactose functionalized 2 nm spherical AuNPs, amine functionalized polystyrene 57 nm spherical nanoparticles, and cationic polysaccharides based on chitosan, are internalized by endothelial cells expressing an intact glycocalyx [18, 22, 25, 26]. We demonstrate that the maturity of the glycocalyx influences the cytotoxic threshold of AuNP+. Cells with a mature glycocalyx can tolerate an 8-fold higher dose of AuNP+ compared to cells with lower glycocalyx expression. This discovery could help explain the observed differences in nanoparticle toxicity between *in vitro* and *in vivo* studies [5, 27, 28].

The exclusion of AuNP-by endothelial cells contrasts with numerous studies that report the internalization of anionic nanoparticles, including AuNPs, across various cell types [29-31]. This discrepancy may partly arise from differences in the physicochemical properties of the nanoparticles, including size, shape, hydrophobicity, net surface charge, and rigidity [32]. However, it is also likely influenced by the state of cells used in *in vitro* assays. Notably, we and others have shown that primary endothelial cells require an extended culture time to produce a mature glycocalyx. This time course of glycocalyx maturity does not correlate with VE-cadherin containing cell-cell junction formation, a condition which can be achieved by tuning the cell seeding density [18, 25, 33]. Many studies expose endothelial cells to nanomaterials only a few hours to a few days after passage, or until confluence is reached, but without specifying the time between passage [34-37].

In this study, we explored how glycocalyx maturity affects interactions with AuNPs. We found that the expression of key glycan and protein components of the glycocalyx increases with culture time. Importantly, when glycocalyx expression is low, AuNP-associate with endothelial cells, but when the glycocalyx is more fully expressed, termed a mature glycocalyx, AuNP-do not. Interestingly, this effect is not observed for polyE, an anionic polymer, suggesting distinct interaction mechanisms for endothelial cells with nanoparticles compared to molecular polymers. In contrast to AuNP-, the level of association of AuNP+ or polyR with endothelial cells remains consistent, regardless of glycocalyx maturity.

Studies involving the selective removal of glycocalyx components provide insights into how engineered nanomaterials interact with the glycocalyx as well as insight into these interactions in various disease states [8, 21]. In this study, using molecular polymers, we show that enzymatic degradation of glycans in the glycocalyx with a combination of hyaluronidase, heparinase III, and neuraminidase moderately increases the association of polyE with endothelial cells, while having no effect on polyR. This supports the idea that the electrostatic interactions between charged polymers and the glycocalyx play a key role, with anionic glycans in the glycocalyx preventing anionic polymers from binding to cells [38]. This finding is consistent with previous reports showing low association of anionic carboxylate functionalized polystyrene spherical nanoparticles (51 nm) with endothelial cells [25].

However, our observation that polyR association with cells is unaffected by glycan presence contrasts with reports showing that cationic amine functionalized polystyrene nanoparticles (57 nm) exhibit increased association with endothelial cells following treatment with hyaluronidase, heparinase III, and neuraminidase [25].Similarly, PEG functionalized AuNPs (10 nm) demonstrate increased intracellular accumulation in endothelial cells treated with heparinase III [16]. In contrast, HS has been reported to inhibit the uptake of AuNPs functionalized with a cationic alkane thiol in the HeLa epithelial cell line [39]. These differences between various engineered nanomaterials, including nanoparticles and molecular polymers, indicate that the glycocalyx serves as a complex barrier whose interactions with engineered nanomaterials are influenced by multiple factors, including the specific glycans present in the glycocalyx as well as the physicochemical properties of the nanomaterials beyond simply surface charge.

## 5. CONCLUSION

This study advances our understanding of how charged nanoparticles and molecular polymers interact with endothelial cells expressing a glycocalyx. Notably, we demonstrate that cationic AuNPs associate with endothelial cells with a mature glycocalyx, whereas anionic AuNPs do not. Additionally, we show that when glycocalyx expression is low, anionic AuNPs associate with endothelial cells, but this effect is not observed for the anionic polymer, polyE. Furthermore, the maturity of the glycocalyx plays a crucial role in modulating the toxicity of AuNP+ with cells containing a mature glycocalyx able to tolerate much higher AuNP+ doses without cytotoxicity than cells with low glycocalyx expression. This suggests that nanoparticles and molecular polymers interact with the glycocalyx through different mechanisms, further influences by the maturity of the glycocalyx. Overall, the findings highlight that while electrostatic interactions are a key factor in engineered nanomaterial-glycocalyx interactions, other aspects, such as the structural properties of both the engineered nanomaterials and the glycocalyx, must also be considered in future research to optimize engineered nanomaterials for biomedical applications.

## Acknowledgements

This work was funded in part by the Australian Research Council (LP190100103 (SMB and MSL) and FT220100092 (MSL)). Confocal microscopy was performed at the Katherina Gaus Light Microscopy Facility and QCM-D performed at the Molecular Surface Interaction Laboratory which are part of the Mark Wainwright Analytical Centre at UNSW and part-funded by the Research Infrastructure program at UNSW.

## Supplementary materials

Supplementary material associated with this article is available.

## References

[1] W. Poon, B.R. Kingston, B. Ouyang, W. Ngo, W.C.W. Chan, A framework for designing delivery systems, Nat Nanotechnol 15(10) (2020) 819–829.

[2] S. Malik, K. Muhammad, Y. Waheed, Nanotechnology: A revolution in modern industry, Molecules 28(2) (2023) 661.

[3] M.J. Mitchell, M.M. Billingsley, R.M. Haley, M.E. Wechsler, N.A. Peppas, R. Langer, Engineering precision nanoparticles for drug delivery, Nat Rev Drug Discov 20(2) (2021) 101–124.

[4] S. Behzadi, V. Serpooshan, W. Tao, M.A. Hamaly, M.Y. Alkawareek, E.C. Dreaden, D. Brown, A.M. Alkilany, O.C. Farokhzad, M. Mahmoudi, Cellular uptake of nanoparticles: Journey inside the cell, Chem Soc Rev 46(14) (2017) 4218–4244.

[5] S. Berger, M. Berger, C. Bantz, M. Maskos, E. Wagner, Performance of nanoparticles for biomedical applications: The in vitro/in vivo discrepancy, Biophys Rev 3(1) (2022) 011303.

[6] D. Chenthamara, S. Subramaniam, S.G. Ramakrishnan, S. Krishnaswamy, M.M. Essa, F.H. Lin, M.W. Qoronfleh, Therapeutic efficacy of nanoparticles and routes of administration, Biomater Res 23 (2019) 20.

[7] J.E. Deanfield, J.P. Halcox, T.J. Rabelink, Endothelial function and dysfunction, Circulation 115(10) (2007) 1285–1295.

[8] S. Weinbaum, J.M. Tarbell, E.R. Damiano, The structure and function of the endothelial glycocalyx layer, Annu Rev Biomed Eng 9 (2007) 121–67.

[9] L. Fu, H.N. Kim, J.D. Sterling, S.M. Baker, M.S. Lord, The role of the cell surface glycocalyx in drug delivery to and through the endothelium, Adv Drug Deliv Rev 184 (2022) 114195.

[10] D.A. Maldonado-Ortega, G. Martínez-Castañón, G. Palestino, G. Navarro-Tovar, C. Gonzalez, Two methods of AuNPs synthesis induce differential vascular effects. The role of the endothelial glycocalyx, Front Med 9 (2022) 889952.

[11] C.S. Alphonsus, R.N. Rodseth, The endothelial glycocalyx: A review of the vascular barrier, Anaesthesia 69(7) (2014) 777–84.

[12] J. Angulo, J. Zimmer, A. Imberty, J.H. Prestegard, Structural biology of glycan recognition, in: A. Varki, R.D. Cummings, J.D. Esko, P. Stanley, G.W. Hart, M. Aebi, D. Mohnen, T. Kinoshita, N.H. Packer, J.H. Prestegard, R.L. Schnaar, P.H. Seeberger (Eds.), Essentials of Glycobiology, Cold Spring Harbor Laboratory Press, Cold Spring Harbor (NY), 2022, pp. 403–18.

[13] A. Aliyandi, C. Reker-Smit, I.S. Zuhorn, A. Salvati, Cell surface biotinylation to identify the receptors involved in nanoparticle uptake into endothelial cells, Acta Biomater 155 (2023) 507–520.

[14] Y.-C. Yeh, B. Creran, V.M. Rotello, Gold nanoparticles: Preparation, properties, and applications in bionanotechnology, Nanoscale 4(6) (2012) 1871–1880.

[15] M.J. Cheng, N.N. Bal, P. Prabakaran, R. Kumar, T.J. Webster, S. Sridhar, E.E. Ebong, Ultrasmall gold nanorods: Synthesis and glycocalyx-related permeability in human endothelial cells, Int J Nanomedicine 14 (2019) 319–333.

[16] M.J. Cheng, R. Kumar, S. Sridhar, T.J. Webster, E.E. Ebong, Endothelial glycocalyx conditions influence nanoparticle uptake for passive targeting, Int J Nanomedicine 11 (2016) 3305–15.

[17] M.J. Cheng, R. Mitra, C.C. Okorafor, A.A. Nersesyan, I.C. Harding, N.N. Bal, R. Kumar, H. Jo, S. Sridhar, E.E. Ebong, Targeted intravenous nanoparticle delivery: Role of flow and endothelial glycocalyx integrity, Ann Biomed Eng 48(7) (2020) 1941–1954.

[18] L. Fu, C.A. Bridges, H.N. Kim, C. Ding, N.C. Bao Hou, J. Yeow, S. Fok, A. Macmillan, J.D. Sterling, S.M. Baker, M.S. Lord, Cationic polysaccharides bind to the endothelial cell surface extracellular matrix involving heparan sulfate, Biomacromolecules 25(6) (2024) 3850–3862.

[19] C.J. Mu, D.A. Lavan, R.S. Langer, B.R. Zetter, Self-assembled gold nanoparticle molecular probes for detecting proteolytic activity in vivo, ACS Nano 4(3) (2010) 1511–20.

[20] Y. Wu, M.R.K. Ali, K. Dansby, M.A. El-Sayed, Improving the flow cytometry-based detection of the cellular uptake of gold nanoparticles, Anal Chem 91(22) (2019) 14261–14267.

[21] J. Joffre, J. Hellman, C. Ince, H. Ait-Oufella, Endothelial responses in sepsis, Am J Respir Crit Care Med 202(3) (2020) 361–370.

[22] R. Gromnicova, M. Kaya, I.A. Romero, P. Williams, S. Satchell, B. Sharrack, D. Male, Transport of gold nanoparticles by vascular endothelium from different human tissues, PLoS One 11(8) (2016) e0161610.

[23] P.H. Olivieri, M.B. Jesus, H.B. Nader, G.Z. Justo, A.A. Sousa, Cell-surface glycosaminoglycans regulate the cellular uptake of charged polystyrene nanoparticles, Nanoscale 14(19) (2022) 7350–7363.

[24] D. Montizaan, R. Bartucci, C. Reker-Smit, S. de Weerd, C. Åberg, V. Guryev, D.C.J. Spierings, A. Salvati, Genome-wide forward genetic screening to identify receptors and proteins mediating nanoparticle uptake and intracellular processing, Nat Nanotechnol 19(7) (2024) 1022–1031.

[25] L. Möckl, S. Hirn, A.A. Torrano, B. Uhl, C. Bräuchle, F. Krombach, The glycocalyx regulates the uptake of nanoparticles by human endothelial cells in vitro, Nanomedicine 12(3) (2017) 207–217.

[26] K.M. Giantsos, P. Kopeckova, R.O. Dull, The use of an endothelium-targeted cationic copolymer to enhance the barrier function of lung capillary endothelial monolayers, Biomaterials 30(29) (2009) 5885–5891.

[27] T. Arokia Femina, V. Barghavi, K. Archana, N.G. Swethaa, R. Maddaly, Non-uniformity in in vitro drug-induced cytotoxicity as evidenced by differences in IC<sub>50 </sub>values – implications and way forward, J Pharmacol Toxicol Methods 119 (2023) 107238.

[28] L. Fu, R. Li, J.M. Whitelock, M.S. Lord, Tuning the intentional corona of cerium oxide nanoparticles to promote angiogenesis via fibroblast growth factor 2 signalling, Regen Biomater 9 (2022).

[29] C. Wilhelm, F. Gazeau, J. Roger, J.N. Pons, J.C. Bacri, Interaction of anionic superparamagnetic nanoparticles with cells: Kinetic analyses of membrane adsorption and subsequent internalization, Langmuir 18(21) (2002) 8148–8155.

[30] E.C. Cho, J. Xie, P.A. Wurm, Y. Xia, Understanding the role of surface charges in cellular adsorption versus internalization by selectively removing gold nanoparticles on the cell surface with a I2/KI etchant, Nano Lett 9(3) (2009) 1080–1084.

[31] S. Mazumdar, D. Chitkara, A. Mittal, Exploration and insights into the cellular internalization and intracellular fate of amphiphilic polymeric nanocarriers, Acta Pharm Sin B 11(4) (2021) 903–924.

[32] R. Augustine, A. Hasan, R. Primavera, R.J. Wilson, A.S. Thakor, B.D. Kevadiya, Cellular uptake and retention of nanoparticles: Insights on particle properties and interaction with cellular components, Mater Today Commun 25 (2020) 101692.

[33] E.M.J. Siren, H.D. Luo, S. Bajaj, J. MacKenzie, M. Daneshi, D.M. Martinez, E.M. Conway, K.C. Cheung, J.N. Kizhakkedathu, An improved in vitro model for studying the structural and functional properties of the endothelial glycocalyx in arteries, capillaries and veins, FASEB J 35(6) (2021) e21643.

[34] A. Aliyandi, S. Satchell, R.E. Unger, B. Bartosch, R. Parent, I.S. Zuhorn, A. Salvati, Effect of endothelial cell heterogeneity on nanoparticle uptake, Int J Pharmaceutics 587 (2020) 119699.

[35] J. Davda, V. Labhasetwar, Characterization of nanoparticle uptake by endothelial cells, Int J Pharmaceutics 233(1) (2002) 51–59.

[36] H. Klingberg, L. B. Oddershede, K. Loeschner, E.H. Larsen, S. Loft, P. Møller, Uptake of gold nanoparticles in primary human endothelial cells, Toxicol Res 4(3) (2015) 655–666.

[37] J. Voigt, J. Christensen, V.P. Shastri, Differential uptake of nanoparticles by endothelial cells through polyelectrolytes with affinity for caveolae, Proc Natl Acad Sci 111(8) (2014) 2942–2947.

[38] M. Wenzel, J. Steup, K. Ohto, J.J. Weigand, Recent advances in guanidinium salt based receptors and functionalized materials for the recognition of anions, Chem Lett 51(1) (2021) 20–29.

[39] B. Peter, N. Kanyo, K.D. Kovacs, V. Kovács, I. Szekacs, B. Pécz, K. Molnár, H. Nakanishi, I. Lagzi, R. Horvath, Glycocalyx components detune the cellular uptake of gold nanoparticles in a size- and charge-dependent manner, ACS Appl Bio Mater 6(1) (2023) 64–73.

